# Unicellular Cyanobacteria Exhibit Light-Driven, Oxygen-Tolerant, Constitutive Nitrogenase Activity Under Continuous Illumination

**DOI:** 10.1101/619353

**Authors:** James Young, Liping Gu, Michael Hildreth, Ruanbao Zhou

## Abstract

Cyanobacteria have played a profound role in shaping the biosphere, most notably through the Great Oxygenation Event (GOE) with the advent of photosynthesis^1^. Cyanobacteria also contribute to global primary production through biological nitrogen fixation (BNF) using nitrogenase^2,3^, an oxygen-labile enzyme complex that evolutionarily predates the GOE^4^. Current literature reports nitrogenase activity in unicellular cyanobacteria is protected from oxygen through diurnal separation of photosynthesis and BNF^5^. However, historic conditions of continuous-light and warm temperature at polar latitudes during the Triassic and Cretaceous may have created a selective advantage amongst unicellular cyanobacteria for non-temporal mechanisms of maintaining nitrogenase activity in the presence of oxygen. Here we report constitutive nitrogenase activity concurrent with a net-gain of oxygen through photosynthesis in a continuous-light adapted culture of the unicellular cyanobacteria, *Cyanothece* sp. ATCC 51142. Nitrogenase activity in the adapted culture exhibited dependence on light and an increased resilience to artificially raised oxygen-tension compared to traditional culture. We predict cyanobacteria closely related to *Cyanothece* sp. ATCC 51142 also possess this physiology and found an accessory predicted proteome with functional relevance. This work provides a model of light-driven, oxygen-tolerant, constitutive nitrogenase activity and suggests this physiology may be conserved in closely related unicellular diazotrophic cyanobacteria with implications for primary production in polar ecosystems and potential biotechnological application in sustainable agriculture production.

## Introduction

Cyanobacteria are reported to overcome the oxygen sensitivity of nitrogenase by employing spatial or temporal mechanisms to create aerobic, micro-oxic, or anaerobic environments^6^. For instance, filamentous *Anabaena* sp. PCC 7120 forms heterocysts to protect nitrogenase in a micro-oxic environment that lacks oxygen-producing photosystem II and performs high respiration^7^. The unicellular cyanobacterium, *Cyanothece* sp. ATCC 51142, was previously reported to separate the two processes of oxygen-producing photosynthesis and oxygen-labile nitrogen fixation temporally within the same cell using a diurnal rhythm^5^. Other unicellular cyanobacteria, like *Cyanothece* sp. PCC 7425 (formerly *Synechococcus* sp. PCC 7425), are dependent on an externally controlled anaerobic environment to fix nitrogen.^8^

Currently reported mechanisms of nitrogen fixation are useful under the prevailing environmental constraints of cyanobacterial growth in dark/light cycles in waters at temperate latitudes. However, recent discoveries of free-living and symbiotic unicellular cyanobacteria in polar regions exposed to continuous-light^9,10^ raises the possibility of non-temporal mechanisms of nitrogenase protection in unicellular cyanobacteria. While filamentous cyanobacteria can fix nitrogen in continuous light within the protective heterocyst, there may be a selective advantage amongst diazotrophic unicellular cyanobacteria capable of non-temporal protection of nitrogenase when exposed to continuous light.

In addition to fixing nitrogen, cyanobacterial nitrogenase also produces hydrogen gas, resulting in overlapping research between these two areas. Recent experiments maintained under a micro-oxic atmosphere suggest a link between photosystems and nitrogenase-catalyzed hydrogen production when a *Cyanothece* sp. ATCC 51142 culture was transitioned from nitrogen-replete (CL N+) to nitrogen-deplete conditions under continuous-light (CL N-)^11^. *Cyanothece* sp. ATCC 51142 was also reported to have high hydrogen production under light when transitioned from the dark phase of a dark-light cycle, concomitant with a net loss of oxygen in both media and headspace when the experiment was run in a closed, non-sparged system^12^. While these experiments focus on hydrogen production under micro-oxic conditions, they hint that unicellular cyanobacteria may be able to harness light to drive nitrogenase activity, consistent with our hypothesis.

## Results

We found sub-culturing *Cyanothece* sp. ATCC 51142 in ASPII N- media in continuous light for many generations (CL N-) resulted in constitutive nitrogenase activity under normal atmospheric conditions. Previous studies with *Cyanothece* sp. ATCC 51142 in CL N- under normal atmospheric conditions only allowed short periods of adaptation, or did not take time-series measurements, likely explaining why this physiology has yet to be reported for *Cyanothece* sp. ATCC 51142.

Following the discovery of constitutive nitrogenase activity, we took oxygraphy measurements of the adapted culture, demonstrating a net gain of oxygen concurrent with nitrogenase activity in our CL N- adapted culture (Figure 1B). While previous reports have shown nitrogenase activity in a non-adapted culture of CL N- *Cyanothece* sp. ATCC 51142, the cultures were still entrained in a cycle, resulting in separation of photosynthesis from nitrogen fixation. We separately validated the oxygraphy data using a secondary experiment measuring nitrogenase activity and oxygen changes over time in non-sparged, closed vials (data not shown here).

**Figure 1:**
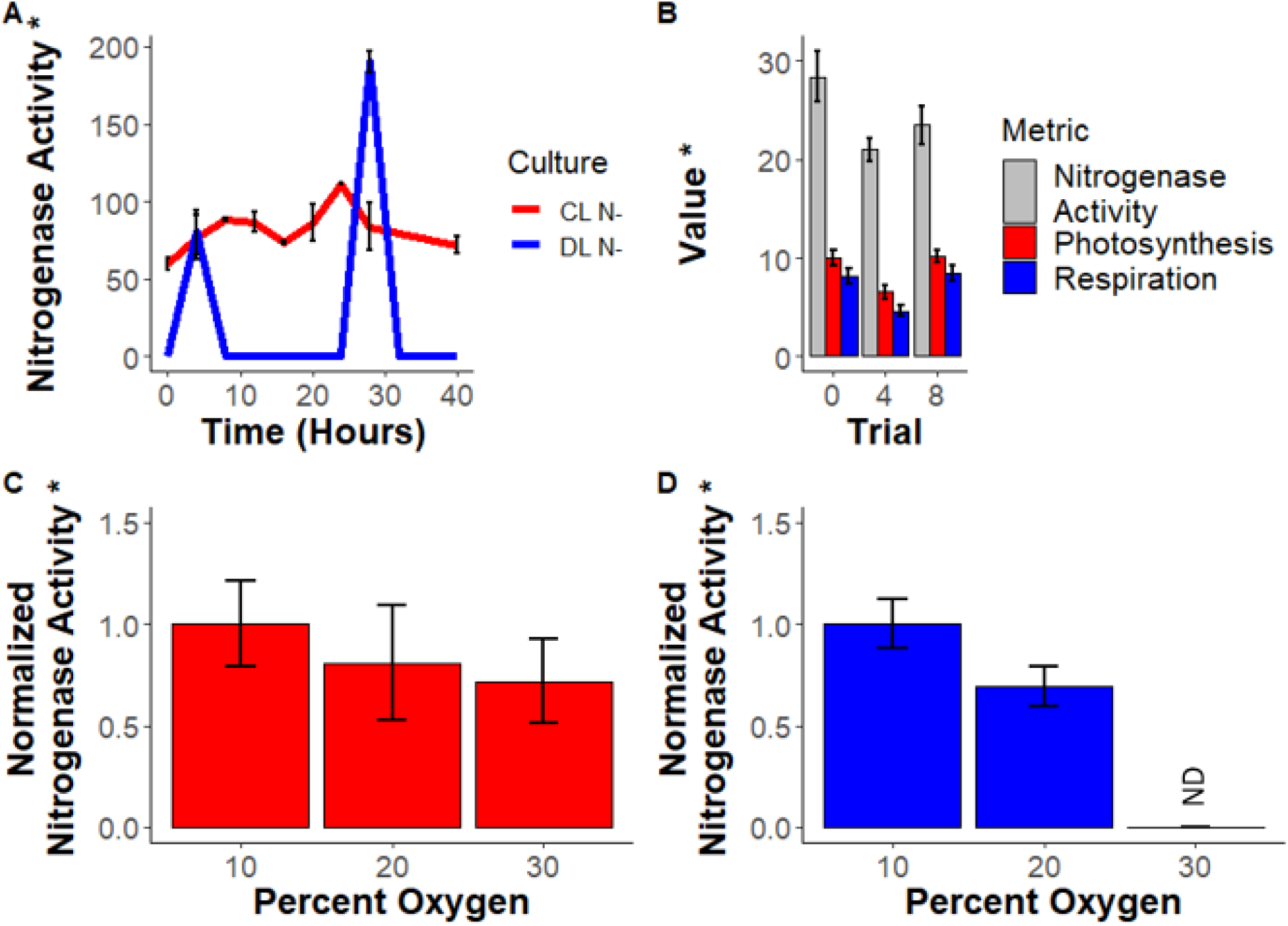
Nitrogenase activity is constitutive and concurrent with oxygenic photosynthesis in adapted CL N- culture (1A, 1B). Nitrogenase activity in the CL N- culture (1C) is more resilient in increasing oxygen concentration compared to DL N- culture (1D). * = Photosynthesis and respiration (1B) units are nmol O_2_ ∙ (mL∙ μg ChlA ∙ min)-1 and acetylene reduction assay units are (integrated units ∙ 10^−6^) ∙ (μg ChlA ∙ mL∙ hour)^−1^, activity in 1A is measured in (nmol ethylene · 10^−6^) ∙ (OD_720_ ∙ mL∙ hour)^−1^. ND = Not Detected. Bars are standard deviation based on n=3.

Next, we aimed to determine the effect of oxygen on nitrogenase activity in this adapted culture. Increasing oxygen produced a small negative effect on nitrogenase activity in the adapted CL N- culture relative to the DL N- culture (Figure 1C and 1D, respectively). This suggests, as expected, oxygen increase in the oxygraphy and headspace experiment were not causal for increased nitrogenase activity, but more likely an artefact of the increased photosystem activity that generates the reductive energy and ATP for nitrogenase activity. This also shows that the CL N- culture has an increased tolerance to oxygen relative to the DL N- culture.

The building experimental evidence suggests a photosystem driven nitrogenase activity, leading us to compare the effect of light on nitrogenase activity between CL N- and DL N- cultures. Our CL N- culture exhibited nitrogenase activity in illuminated incubation while the dark incubated CL N- culture had undetectable nitrogenase activity (Figure 2C) which is indicative that this physiology is photosystem driven. DL N- culture tested during peak nitrogenase activity exhibited nitrogenase activity in both illuminated and dark incubation, with slightly higher activity in under illumination (Figure 2D), possibly due to photo-cyclic phosphorylation while PSII was shut-down by the PsbA sentinel protein^13^.

**Figure 2:**
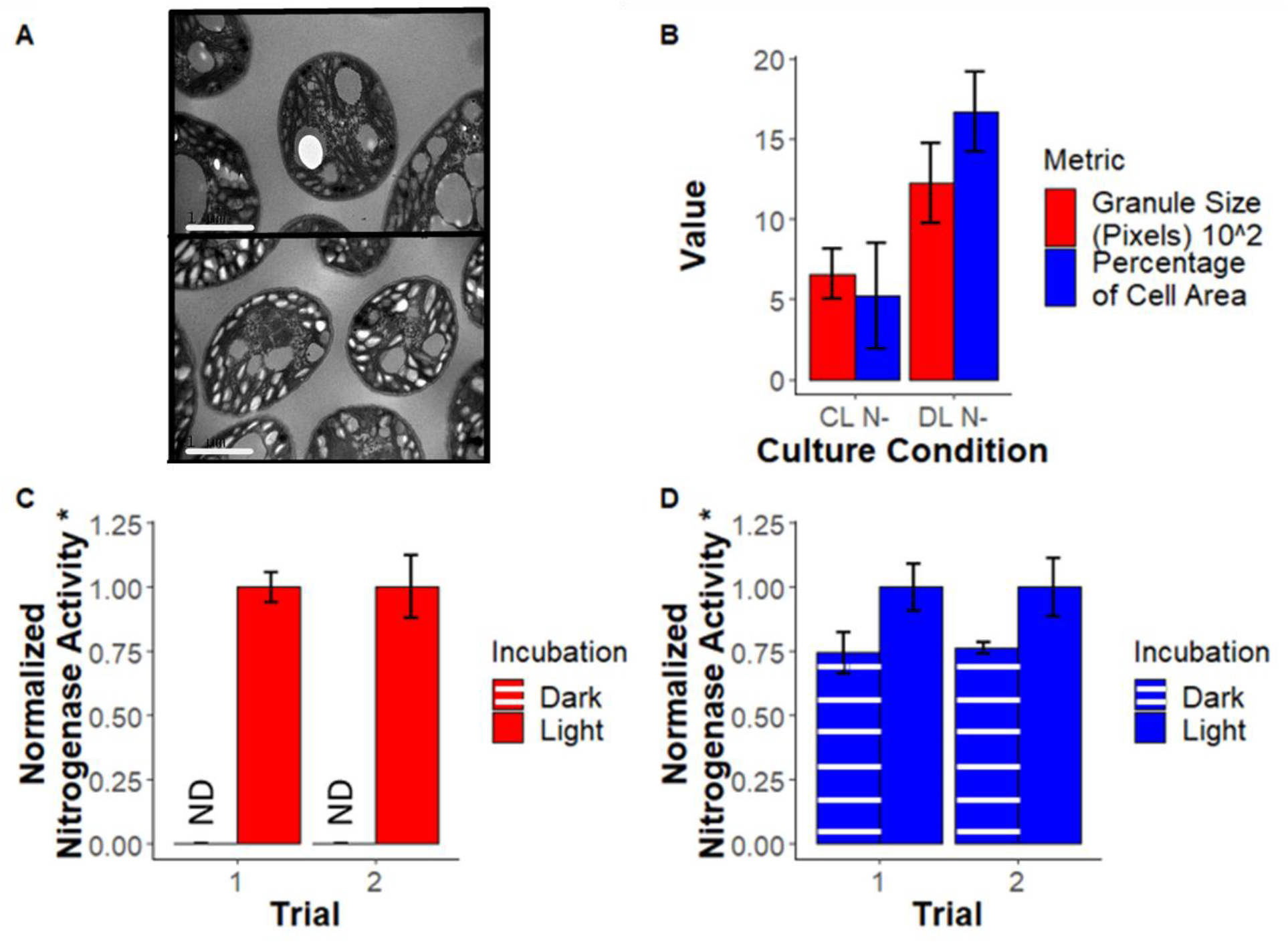
Bacterial glycogen granules (smaller white granules toward cell periphery) occupy less space in cells from CL N- culture (2A top) than DL N- culture (2A bottom) with the difference being quantified using ImageJ (2B). White bars represent 1μm (2A). CL N- culture nitrogenase activity is dependent on illumination (2C) while DL N- exhibits nitrogenase activity regardless of illumination (2D). Acetylene reduction units are (integrated units ∙ 10^−6^) ∙ (μg ChlA ∙ mL∙ hour)^−1^. ND = Not Detected. Bars for 2C and 2D are standard deviation based on n=3. Bars for 2B are standard deviation based on cells analyzed (CL N- n=5, DL N- n=6)

In addition to undetectable nitrogenase activity in the dark, CL N- cells had less bacterial glycogen granules than DL N-, as observed by TEM imaging (Figure 2A and 2B). The DL N- culture presented many bacterial glycogen granules, consistent with previous literature^14^. Bacterial glycogen granules are produced and stored during the day to fuel the respiration necessary for dark constrained nitrogen fixation in DL N- cultured *Cyanothece* sp. ATCC 51142. The lower presence of bacterial glycogen granules in the CL N- culture adds to the evidence suggesting this adapted CL N- physiology is more dependent on photosynthesis than respiration for nitrogenase activity. However, the oxygraphy tests show respiration still occurs at high levels relative to the CL N+ culture, indicating that relatively high respiration is still associated with this physiology (data not shown here).

After discovering oxygen-tolerant nitrogenase activity in *Cyanothece* sp. ATCC 51142, we further investigated the literature for closely related cyanobacteria that might suggest conservation of this physiology. One paper reported similar physiology in *Gloeothece* sp. 68DGA^15^. Further supporting our prediction of physiological conservation, *Crocasphaera watsonii* WH8501 has been reported to grow diazotrophically under continuous light for weeks^16^. However, the data was not made available and the relation of photosynthesis and nitrogen fixation was not investigated. *Cyanothece* sp. PCC 8801, formerly *Synechococcus* sp. RF-1, has also been reported to perform continuous nitrogenase activity in continuous light, but its photosynthetic regime was not investigated^17^. We found the cyanobacteria predicted to share this physiology share a most recent common ancestor that is absent from other *Cyanothece*, such as *Cyanothece* sp. PCC 7425^18^. This information indicates that *Gloeothece* sp. 68DGA, *Cyanothece* sp. ATCC 51142, *Cyanothece* sp. PCC 8801, and *Crocasphaera watsonii* WH8501 may all share this ability, despite a current lack of data.

Supporting this prediction, analysis of UniProt predicted proteomes revealed a unique accessory proteome common between *Cyanothece* sp. ATCC 51142, *Crocasphaera watsonii* WH8501, and *Cyanothece* sp. PCC 8801 (CL N- group) but absent from the control group. The control group consisted of the pan-proteome constructed from the UniProt predicted proteome of anaerobic diazotroph *Cyanothece* sp. PCC 7425 and closely related but non-diazotrophic *Synechocystis* sp. PCC 6803. These comparisons were made with *Cyanothece* sp. ATCC 51142 as the reference database with a 50% amino acid sequence similarity as the cutoff. We found 431 unique proteins in the CL N- group and 686 predicted proteins unique to the control group. To gain an understanding of the functional significance of these predicted proteins we used public data from a recent transcriptome project^19^ to analyze differential correlation of these genes between DL N- and CL N- conditions over time. While the transcriptome project did not observe the physiology we are reporting, likely due to the short transition time from DL N-, the start of the adaptive response to continuous light still gives valuable insights that can be expounded on in later experiments. We followed the differential correlation analysis with a network analysis and GO enrichment analysis to allow functional inferences.

The accessory proteome of the CL N- group had higher network inter-connectivity compared to the control (Figure 3A and 3B, respectively) indicating these genes are more orchestrated in expression changes between dark/light cycled and continuous light conditions. The CL N- group had significantly higher GO enrichment for transmembrane transport and cellular metabolic processes (Figure 3C). Genes unique to the CL N- group with significant differential correlation (p<0.001) between DL N- and CL N- were investigated for functional implications. A three-gene operon (cce_0574-cce_0576) located near the nif-operon is predicted to facilitate ferrous ion uptake. A predicted NifU-like gene (cce_1857) was also found to be unique to the CL N- group. Iron is an important part of the nitrogenase Fe protein and NifU proteins are involved in Fe-S cluster construction and repair. Other genes of interest are predicted to be involved in transcription and translation regulation or protein-protein interactions. Most predicted genes found to be significantly differentially expressed were hypothetical proteins, many of which were predicted to be membrane proteins.

**Figure 3:**
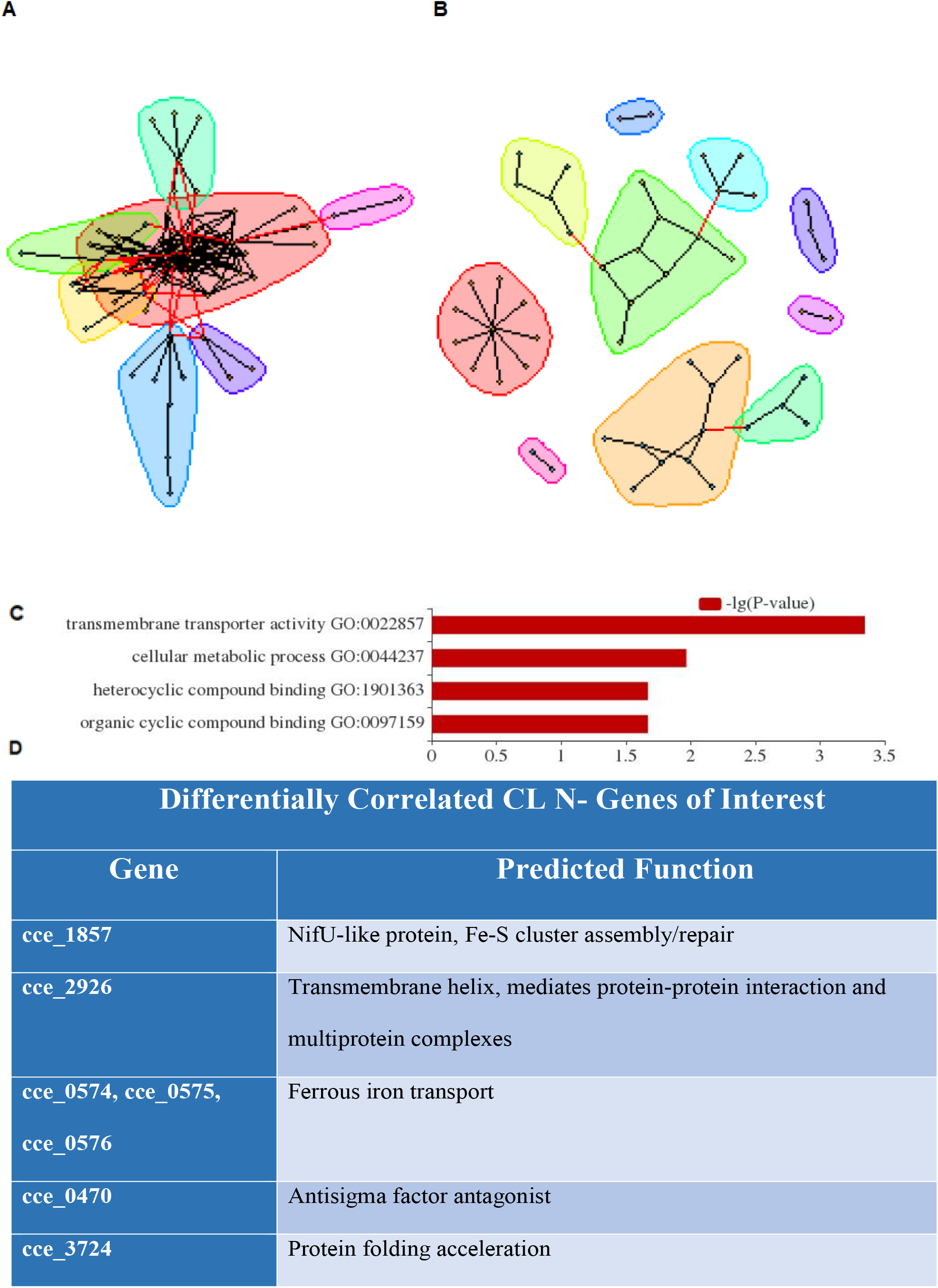
The genes with differential correlations between DL N- and CL N- with adjusted p- values <0.001, the predicted accessory proteome had higher connectivity (3A) while the control group had lower connectivity (3B). The predicted accessory proteome with differential correlation with adjusted p-value < 0.001 was enriched for various gene ontologies compared to the control group (3C). Genes of interest were found based on predicted function that could play a role in our observed physiology (3D).

## Discussion

This work establishes a model culture for oxygen-tolerant nitrogenase activity as evidenced by comparative measurements of CL N- and DL N- nitrogenase activity with complementary oxygraphy, headspace oxygen, and manipulated oxygen concentration experiments. We began elucidation of the mechanisms through acetylene reduction assays and TEM ultrastructure analysis. We predict this adaptive ability may be conserved in cyanobacteria originating from a shared common ancestor. The foremost question we interrogate below is if nitrogen fixation and photosynthesis are occurring simultaneously in single cells. Regardless of the culture’s tendency towards physiologic homogeneity or heterogeneity, there is nitrogenase activity concurrent with a net gain of oxygen, suggesting some mechanism of oxygen-tolerance. We also discuss the possible conservation of this ability and its implications for ecosystem function and biotechnological application.

There have been limited previous attempts to determine if an individual cell can fix nitrogen and produce oxygen simultaneously, likely due to the counter-intuitive nature of the proposition. *Gloeothece* sp. 68DGA is reported to have a homogenous and time-tolerant distribution of NifH protein throughout the culture when adapted to continuous light, suggesting all cells are fixing nitrogen^15^. One of the few papers to-date citing the *Gloeothece* sp. 68DGA paper observed *Crocasphaera watsonii* WH 8501 and determined individual cells could not fix carbon and nitrogen simultaneously to a large extent^20^. However, this *Crocasphaera watsonii* WH 8501 culture was only exposed to continuous light for 24 hours, the acetylene reduction assays indicated the culture was still entrained in a rhythm, and oxygen production was not measured.

We propose two explanations for our observed physiology. The first possible explanation is that the culture is physiologically heterogenous where a portion of the cells are performing high respiration to protect nitrogenase from oxygen and provide reducing equivalents while the other portion of the culture is performing photosynthesis. Our second explanation is that the culture is homogenous and most of the cells are fixing nitrogen and performing photosynthesis simultaneously. Our results, when evaluated together, are not consistent with physiological heterogeneity and indicate the higher likelihood of a physiologically homogenous culture.

If individual cells in the CL N- culture were relying solely on high respiration to protect and support nitrogenase activity, they should still exhibit nitrogenase activity when incubated in the dark. However, we observed undetectable levels of nitrogenase activity in this scenario (Figure 2C) indicating the necessity of light and implicating photosystems as a driving force for this physiology. We also observed a net gain of oxygen concurrent with nitrogenase activity in the CL N- adapted culture (Figure 1B), indicating that a heterogenous culture would need individual cells with respiration rates high enough to decrease their local oxygen without decreasing the total oxygen. This seems unlikely, especially considering the culture is planktonic and shaken. The lower content of bacterial glycogen granules in CL N- relative to DL N- also contradicts the proposition of individual cells with abnormally high respiration in CL N-, assuming the hypothetical “high respiration” cell would fuel respiration with stored carbohydrates in the bacterial glycogen granules consistent with previous literature^21^. The results of this multi-faceted interrogation indicate the higher likelihood of a physiologically homogenous culture than a heterogenous culture.

This physiological adaptation, which we predict is conserved in the CL N- group (Figure 3) may have originated from a selective advantage in response to environmental parameters present in the warm, open waters at polar latitudes that were exposed to continuous light during the Triassic^22^ and Cretaceous^23^ periods. The oceans were also highly nutrient limited in the Jurassic^24^ likely flowing over into the Cretaceous and creating a demand for nitrogen fixation as has been proposed for the symbiotic UCYN-A, which is closely related to *Cyanothece* sp. ATCC 51142^18^. Recent phylogenetic work indicates *Cyanothece* sp. ATCC 51142 and *Crocasphaera watsonii* WH8501 diverged from their most recent common ancestor during the early Triassic and *Cyanothece* sp. ATCC 51142 from *Cyanothece* sp. CCY 0110 occurred near the beginning of the Cretaceous, preceding the divergence of UCYN-A1 and UCYNA-2 from their respective common ancestor^18^.

UCYN-A became endosymbiotic in pelagic prymnesiophytes approximately 91 million years ago^18^. Recent research has found that UCYN-A is actively fixing nitrogen in the Arctic Ocean^10^ and that UCYN-A preferentially expresses nitrogen fixation genes in the light^25^ but it doesn’t produce an oxygen since it has lost PSII throughout coevolution with its symbiotic partners. *Cyanothece*-like free living cyanobacteria from the Chroococcales order, which may share this physiological adaptation, have also been found actively expressing nifH genes in arctic latitudes^9^. With continuing loss of icecaps causing an amplification of surface temperature warming^26^, the growth of continuous light adaptable diazotrophs toward the poles may become increasingly relevant to marine ecosystem function considering their primary production role^27^ (Figure 4).

**Figure 4:**
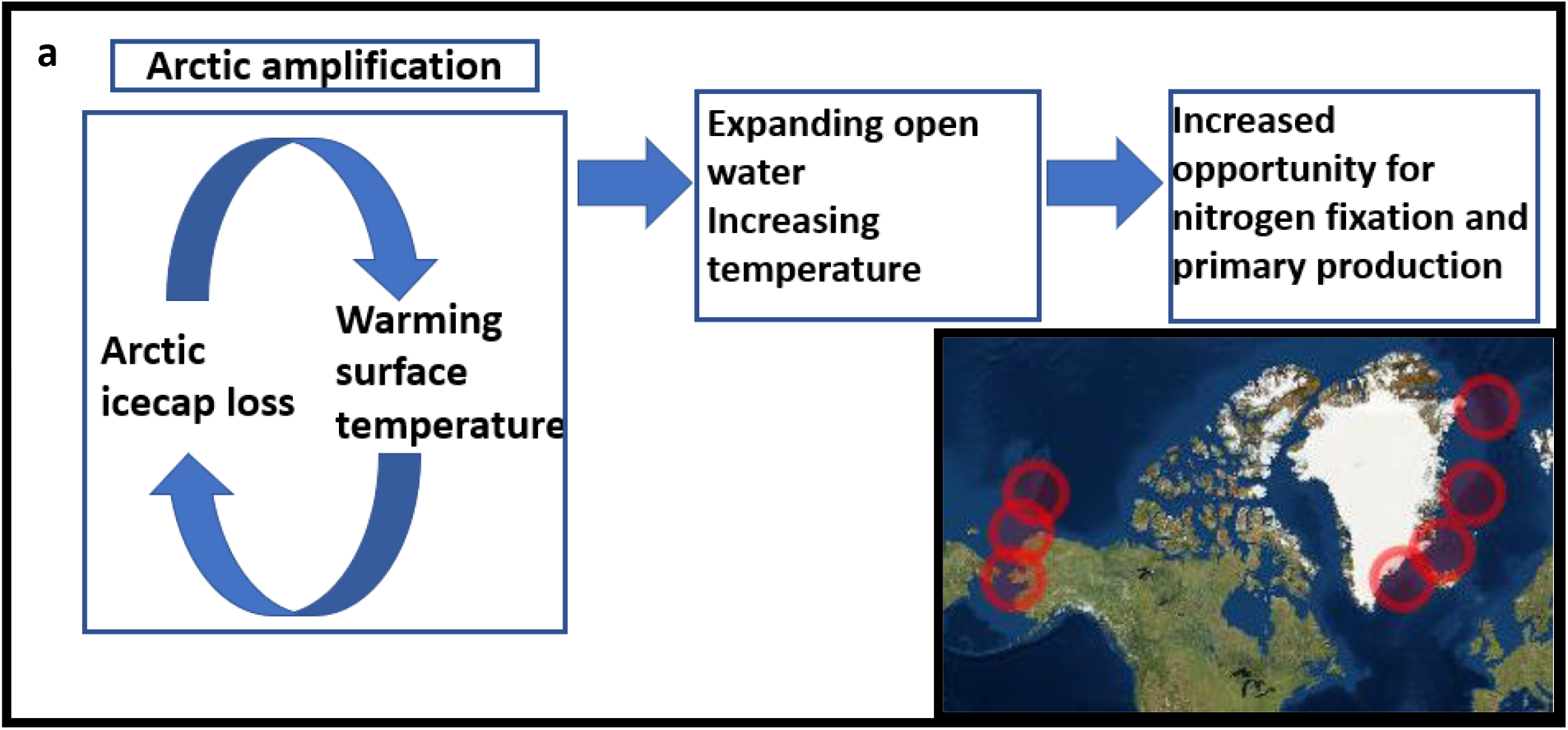
Primary production of unicellular diazotrophic cyanobacteria in the arctic has potential to increase due to increasing open water and rising temperatures at latitudes that experience continuous light for months (4A). Cyanobacteria closely related to Cyanothece have been recently discovered in the Arctic Ocean^9,10^ (4B).

In addition to environmental implications, *Cyanothece* sp. ATCC 51142 provides a model of oxygen-tolerant nitrogenase activity, which upon elucidation can be emulated in attempts at engineering BNF into crops. Recent attempts to transfer a functional complement of nitrogenase genes to *Synechocystis* sp. 6803 have demonstrated the complexities engineering BNF into a non-diazotrophic phototroph^28,29^. The engineered cells exhibited nitrogenase activity at anaerobic levels, but cells exposed to 1% oxygen quickly lost ~90% of activity. The genes identified in our bioinformatic analysis (Figure 3D) mark a starting point for further inquiry into the mechanism of this physiology which may lend further insights to BNF engineering projects.

## Methods

### Culture Conditions

*Cyanothece* sp. ATCC 51142 cultures were cultivated batch style and sub-cultured for at least ten generations in ASPII N- (nitrogen deplete) or ASP II N+ (nitrogen replete) media. The continuous light cultures were grown at 30°C on a circular shaker at 120rpm under continuous μE m^−2^ s^−1^ light. Cultures were refreshed weekly leading up to experiments. The 12-hour dark / 12-hour light cycled cultures were grown in an Innova 44® Incubator set at 30°C on a circular shaker at 120rpm. The light period for the 12-hour dark / 12-hour light cycled culture was grown under GE gro-lights. When preparing culture for acetylene incubation to determine nitrogenase activity, 5 mL’s of culture was transferred to a 20 mL glass tube. The seal used on the glass tube was dependent on the experiment. The continuous nitrogenase activity experiment used a red cap while the headspace oxygen experiment used a blue air-impermeable cap.

### Continuous Nitrogenase Activity Measurements

Five mL of the three to eight-day old cultures growing under continuous light (μE m^−2^ s^−1^., 120 rpm, 30°C) or 12 h light/12 h dark cycle in ASPII N+ or ASP II N- (nitrogen-containing or nitrogen-free media) were transferred into a 20-ml glass serum bottle (Wheaton) respectively. The bottle was sealed with a red rubber stopper (Wheaton) and injected with 0.5 ml of acetylene. The bottle was incubated under light or dark for one hour. Five mL of headspace gas sample from the 20 mL culture bottle was administered via a 1 mL GSV Loop to the GC-MS (Agilent 7890A/5975C). The volatile compounds were separated by CP7348 column (Agilent PoraBOND Q 25 m × 250 µm × 3 µm) with Pulsed Split Mode at 100:1 ratio. The carrier gas was hydrogen at a flow rate of 0.8 mL/min, and the supply of Ultra-pure N_2_ and Ultra-Zero air for the FID were 15 and 200 ml/min respectively. The GC program was initiated at 32°C held for 4 min, and ramped at 110°C to reach 232°C. The temperatures of the valve box heater and flame ionization detector (FID) heater were 100°C and 250°C, respectively. The scanning mass range of MSD was between 10 to 50 *m/z* and the inlet temperature was set at 250°C. All measurements were performed in triplicate.

### Oxygraphy Measurements

Oxygraphy measurements were taken using a Hansa Oxygraph II and Hansa Light Source. The light source was set at μE m^−2^ s^−1^ and the temperature of the chamber was kept constant using a circulating chiller set at 30°C. Cultures were concentrated from the 3 replicate tubes used for each acetylene reduction, 15 mL’s total, down to 3 mL’s total by centrifugation and removing supernatant. Before the concentrated samples were transferred to the oxygraphy chamber, sodium bicarbonate was added and brought to a final concentration of 10 mM. The addition of sodium bicarbonate served to make sure there was an adequate supply of carbon dioxide to the cyanobacteria since there is no headspace in the prepared oxygraphy chamber. Samples came to relative stability (steady rate of respiration) in the dark before the light source was turned on, photosynthesis was measured for at least two minutes or until the rate stabilized, then the light source was turned off and respiration was measured for at least 2 minutes or until stabilized. The stir bar setting was set to 75% and the equilibration of the device was completed with distilled deionized water at 30°C that was sparged by 99% nitrogen gas.

### Simultaneous Measurement of Nitrogenase Activity and Headspace Oxygen

For each experimental replication, all samples were sealed in air impermeable glass tubes (20 mL) at the zero hour with 5 mL of culture. Each tube had 0.5 mL of 99% carbon dioxide added to prevent a loss of photosynthesis over the 12 to 16 hours incubation period. Carbon dioxide was added to all tubes, including the control and zero to one-hour incubation samples for consistency. Tubes had 0.5 mL’s of pure acetylene added immediately preceding their designated 1-hour incubation period. After acetylene incubation, 5 mL’s of headspace were withdrawn for determination of nitrogenase activity (ARA). An additional 5 mL was taken from the same tube and injected into a GC autosampler vial for oxygen content measurement. The GC autosampler vials were degassed preceding the experiment by filling and withdrawing the vials with 80% hydrogen, 20 % carbon dioxide gas to 20 psi. Each vial underwent 6 rounds of 30 seconds gas and degas, ending with a gas fill. The headspace oxygen analysis was performed using GC coupled with thermal conductivity detector (TCD) (Shimadzu GC 14B equipped with a CombiPal AOC-5000 auto-sampler and an MSH 02-00B injector needle with a 2 mL injection loop, SHIMADZU Corp.). The carrier gas was Helium at 275 kPa, and the supply of Ultra-Zero Air, 95% Argon-5% Methane, and H_2_ were 177.8, 167.5, and 510 kPa respectively. The column had a total flow rate of 20 mL/min and a purge flow rate of 1.0 mL/min and a pressure of 323.0 kPa. The temperatures of the injection loop, oven, and TCD were 100, 90, and 100°C respectively. GC-TCD data were obtained by passing sample through four columns in series (Hayesep N 80/100 mesh 1.50 m × 1/8 IN ×2.1 mm SS, Hayesep D 80/100 mesh 2.50 m × 1/8 IN × 2.1 mm SS, Hayesep D 80/100 mesh 2.50 m × 1/8 IN × 2.1 mm SS, and Supelco 60/80 Molecular Sieve 5 Å 3.0 m × 1/8 IN × 2.1 mm SS) for 13-min isothermal period at 90 °C. For quality assurance, all measurements were performed in triplicate.

### Exogenous Oxygen Manipulation

To control the oxygen content, a combination of withdrawing headspace and adding volumes of gas were employed. Tubes were then incubated for an hour in their native environment, light or dark, then sampled for acetylene reduction as mentioned previously. All measurements were performed in triplicate.

### Light Dependent Incubation

Continuous-light and 12-hour dark / 12-hour light cycled culture was added to sealed glass tubes (20 mL) as described above and had 0.5 mL acetylene added. The tubes were incubated either under illumination or the dark with temperature and shaking speed held constant at 30°C and 120 rpm. Headspace was withdrawn and analyzed for acetylene reduction as a representation of nitrogenase activity as described above using the Agilent GCMS.

### TEM microscopy

Triplicate acetylene reduction assays were measured to quantify nitrogenase activity. Twenty mL glass tubes containing five mL culture each were combined and centrifuged at 12,000×g for 10 minutes. Supernatant was removed and the pellet was washed twice in 5mL of 0.1M Millonig’s phosphate buffer, 7.4 pH, centrifuged at 12,000g for 5 minutes. After two washes the pellet was resuspended once more in 1 mL of 0.1M Millonig’s phosphate buffer, 7.4 pH, and transferred to a 1.25 mL capacity, externally threaded, conical bottom, free-standing tube. This tube was centrifuged at 12,000×g for 10 minutes and had the supernatant removed and was then covered with 2.5% glutaraldehyde in 0.1M Millonig’s phosphate buffer, 7.4 pH. The tube was incubated at room temperature for 1 hour then stored at 4°C for less than 6 hours before embedding.

After this time period the samples were washed twice in the buffer, post-fixed with 2% OsO_4_, buffer rinsed twice and refrigerated overnight. The samples were then rinsed twice in water, washed twice each with 30% acetone, 50% acetone, 70% acetone and uranyl acetate, 90% acetone, and 100% acetone. The samples were then embedded in Epon plastic, sectioned using a diamond knife and ultra-microtome, transferred to copper grids, and stained with lead citrate.

Images were captured on JEOL JEM-100CX II tungsten-filament 100kV transmission electron microscope. Digital imaging was conducted with Gatan Erlangshen ES500W camera, using Gatan Digital Micrograph software. ImageJ was used for extracting bacterial glycogen granule size using thresholding set at 0 and between 110 and 120.

### Bioinformatic Analysis

UniProt predicted proteomes were used for selected bacterial strains and compared to *Cyanothece* sp. ATCC 51142 using BLAST+. Proteins were found to be unique to the “CL N-” group accessory proteome or the control group pan-proteome at 50% amino acid sequence similarity to *Cyanothece* sp. ATCC 51142. These predicted proteins were then interrogated at the transcriptome level from a recent experiment^19^ for differential correlation between lighting conditions using DiffCor in R. There were 11 time points taken from the DL N- cycle and 9 time points taken from the CL N- cycle. Gene pairs with highly significant differential correlation (p < 0.001) were visualized as networks using MEGENA in R. We also looked for enrichment of gene ontology annotation to make inferences of functional roles of the genes using UniProt.

## Acknowledgements

This work was partially supported by USDA-NIFA GRANT Heterocyst Transcriptomics (to R. Z.), and by the South Dakota Agricultural Experiment Station.

